# Deep FASTQ and BAM co-compression in Genozip 15

**DOI:** 10.1101/2023.07.07.548069

**Authors:** Divon Mordechai Lan, Daniel S.T. Hughes, Bastien Llamas

## Abstract

We introduce Genozip Deep, a method for losslessly co-compressing FASTQ and BAM files. Benchmarking demonstrates improvements of 75% to 96% versus the already-compressed source files, translating to 2.3X to 6.8X better compression than current state-of-the-art algorithms that compress FASTQ and BAM separately. The Deep method is independent of the underlying FASTQ and BAM compressors, and here we present its implementation in Genozip, an established genomic data compression software.

## Introduction

The Institute of Genomic Medicine’s (IGM) Bioinformatics Core, situated within the Columbia University Irving School of Medicine, manages a variant warehouse containing approximately 130,000 whole-genome sequencing and whole-exome sequencing samples. This warehouse serves the dual purpose of gene discovery and diagnostic analysis. Given that the BAM (Binary sequence Alignment/Map) files used to generate the warehouse have been used as the foundation for numerous publications and diagnostic analyses, and continue to be reanalysed, the IGM is obliged to store these files in their current format for the foreseeable future. Additionally, the IGM acts as a long-term repository for off-machine raw sequencing data (FASTQ files) of internally and externally sequenced samples, which must be preserved in their original form. Currently IGM has around 5 petabytes of storage of which the vast majority are FASTQ files compressed with gzip and BAM/CRAM files. While these file types are already compressed, the rapid growth of the volume of data puts the IGM in dire need of improved compression methods. This situation is far from being anecdotal and is a major concern for many institutions and organisations that rely heavily on genome sequencing to support their biomedical and clinical research agendas.

Several commercial and open source software packages have been introduced in recent years for compressing FASTQ files, and others for compressing BAM files, with a handful capable of compressing both BAM and FASTQ, but separately^1–4^. Looking for a new method to address the needs of IGM and other similar users of Genozip, we decided to focus on the large overlap in information content between a typical BAM file and the set of FASTQ files used to generate it. Here, we present a novel method, Deep, for co-compression of BAM and FASTQ files. Deep exploits the information overlap to improve compression, an approach that has not been attempted before, while still guaranteeing losslessness for both FASTQ and BAM data. We demonstrate that this method results in substantially smaller files than when compressing BAM and FASTQ files separately—resulting in a co-compressed file containing both the BAM and FASTQ data with a size that is only slightly larger than just the BAM file compressed with Genozip.

We implemented the Deep method on top of the existing Genozip platform—an established software package for compressing genomic files^5–7^. We released the resulting combined system as Genozip version 15. The `--deep` command line option triggers lossless Deep co-compression of a BAM file with the set of one or more FASTQ files from which the BAM file originates. Genozip automatically manages decompression and processing of a Deep-compressed file using its standard commands, without further options: `genounzip` reconstructs the entire set of input files (BAM and FASTQ), while `genocat` allows the extraction of a single file (i.e. the BAM file or one of the FASTQ files).

## Methods

The Deep method must be implemented on top of a compressor already capable of compressing BAM and FASTQ data. We have implemented it within the Genozip system, but the method described hereinafter is not specific to Genozip, and could be implemented in other suitable compressors. We shall focus our discussion on describing and analysing the Deep method (see Table S5 for source code information). We refer the interested reader to earlier publications^5,7^ describing the methods Genozip utilises to compress the actual BAM and FASTQ data.

It seems obvious that co-compression of FASTQ and BAM would be beneficial, given that read names, sequences and base quality score strings of related FASTQ and BAM files are expected to be similar. However, there are several hurdles that make directly exploiting this information redundancy challenging—in particular doing so fast enough and with economical enough utilisation of RAM, to make it useful for real-world large institutional deployments. These hurdles include: reads in the BAM file are often ordered differently than in the FASTQ file, since it is common practice to sort BAM files by genomic coordinates. Sometimes, read names differ between the FASTQ and BAM data—we have encountered read names changed to include the FASTQ file identifier, to include a unique molecular identifier, to include the sequence length, to conform with the NCBI SRA read name format, or to be more concise by reduction to a sequential numerical number. The base quality (QUAL) data may differ as well—for example if the BAM data underwent Base Quality Score Recalibration. The BAM file might be missing reads contained in the FASTQ data due to filtering, and conversely may include secondary and supplementary alignments not present in the FASTQ data. The nucleotide sequence (SEQ) data in the BAM file might be reverse-complemented, and the QUAL data reversed, versus the FASTQ strings. SEQ and QUAL strings in the BAM file might be shorter than in the FASTQ file due to trimming or cropping. Finally, it is common to map multiple FASTQ files into a single BAM file.

Our method consists of four modules, as follows (Figure 1).

**Figure 1.**
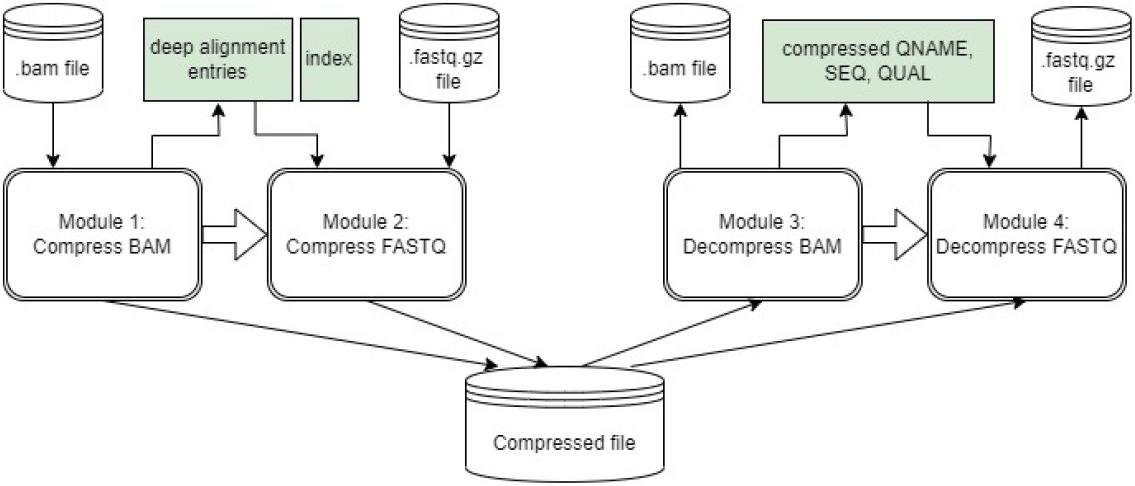
Module 1 is executed during the compression of a BAM (or SAM or CRAM) file, which is compressed first. During the compression process, a “deep alignment entry” comprised of hash values of QNAME, SEQ and QUAL is stored in RAM, and indexed by a value derived from SEQ. Module 2 is run during the compression of FASTQ data: for each read, we use the index to lookup candidate deep alignment entries and determine whether the read is present in the BAM data. If it is, we represent it in the compressed file as a reference to the matching BAM alignment rather than compressing the sequence, base quality and read name data explicitly. Module 3 and 4 are utilised during decompression. Module 3 runs when decompressing the BAM file, compressing and storing in RAM the QNAME, SEQ and QUAL information of each primary alignment. When the FASTQ data is decompressed, Module 4 is deployed to retrieve this information from RAM and reconstruct the FASTQ reads.

Module 1 is run during BAM compression: when compressing each of the BAM alignments, if the alignment is not a supplementary or secondary alignment, Genozip also generates a *deep alignment entry* in RAM corresponding to the alignment. The deep alignment entry consists of 32 bit hash values for each of the QNAME, SEQ and QUAL fields, a *place* field which is the location of the alignment in the BAM file, and a *consumed* flag which is reserved for use in Module 2. In case the *reverse complement* bit of the FLAG field is set, the SEQ string is reverse complemented and the QUAL string is reversed prior to calculating the hash values. In addition to the array of deep alignment entries, Module 1 also generates a *deep index*. The deep index is a hash table, in which each deep index entry contains a linked list of indices into the deep alignment entries array, of all deep alignment entries that are mapped to this particular deep index entry. The deep index entry to which a deep alignment entry is mapped, is determined by a subset of the bits of the *SEQ hash value* of deep alignment entry. The number of bits is a function of the estimated number of alignments in the BAM file.

Module 2 is run during FASTQ compression, and is the most complex of the four modules: at initialisation, this module inspects the first few reads in the FASTQ file, calculating the hash values of the read name, SEQ and QUAL, and looking for matching hash values in the deep alignment entries which were previously stored in RAM by Module 1. Based on whether such matches exist or not, the module determines the Deep mode to be used, which is one of four options: SEQ + read name + QUAL (if all three fields tend to have a match in the BAM data), SEQ + read name, SEQ + QUAL (if only SEQ and either read name or QUAL fields tend to match) or none at all. Then, for each read being compressed, Module 2 does two things: First, it determines whether this read possibly exists in the BAM file based on using the *deep index* stored by Module 1 in RAM to find a *deep alignment entry* with a matching hash value of SEQ and a matching hash value of at least one of read name and QUAL (depending on the Deep Mode). Second, crucially, given a set of hash value matches, Genozip ascertains that the data itself match as well, despite not having access to the BAM data—as we store only the hash values of the read name, SEQ and QUAL it in RAM, not the actual strings. If the Module is certain that this FASTQ read has exactly one matching alignment in the BAM file, then it sets the *consumed* flag in the deep alignment entry, and represents, in the compressed output file, the matching read name, SEQ and/or QUAL data as a reference to the *place* in the BAM file, where *place* is extracted from the deep alignment entry. This representation of the FASTQ read components as a reference to the BAM data, rather than compressing them explicitly, is the crux of how the Deep method improves compression.

The ascertainment that the hash match indeed refers to the BAM alignment derived from the current FASTQ read, but not to another unrelated alignment that by chance has the same hash values, is done as follows: first, the entire linked list in the matching deep index entry is inspected for matching hash values. If more than one deep alignment entry on the linked list has matching hash values, i.e., the current FASTQ read maps to multiple BAM alignments, then we abandon the Deep method for this read, as we don’t know which of the matching BAM alignments corresponds to this FASTQ read, and instead fall back to Genozip’s regular method for compressing a FASTQ read. If there is a single match, but the *consumed* flag in the deep alignment entry has already been set by a prior FASTQ read, this indicates that multiple FASTQ reads map to a single BAM alignment. Because we use a 64 or 96 bit value (32 bits for each of SEQ, QUAL and read name), it is extremely unlikely that two different FASTQ reads will map to the same BAM alignment (one of them incorrectly so). If this does happen, we abandon the compression and advise the user that the `--deep` option cannot be used with these files. To prevent this from happening trivially, we exclude reads with a SEQ that is a string of a single character (N or a base). If we had left it at that, there could still be an edge case where a FASTQ read could have been matched with an incorrect BAM alignment due to chance equivalence of the hash values. This could happen if, for example, there are two FASTQ reads that by chance have the same hash values, where one of these reads does not have a corresponding alignment in the BAM file because it was filtered out, and the other read, which does have a corresponding alignment, is in a FASTQ file that the user omitted from the `genozip` command line. In this case, Genozip might incorrectly determine that there is a unique match between the sole read and the sole alignment with these hash values. To avoid this edge case, Genozip requires that all FASTQ files that contributed reads to the BAM data are provided as inputs. If not all FASTQ files are provided and this edge case does occur, Genozip will catch it during the testing phase that follows the compression, during which Genozip verifies that the compressed data is reconstructable losslessly.

Module 3 is run during BAM decompression: when decompressing a non-supplementary, non-secondary alignment, this module compresses the SEQ, QUAL and QNAME data and stores them in RAM, in an array indexed by *place* (i.e., the sequential number of this alignment in the BAM file). If the alignment has the *reverse complemented* flag set, SEQ is stored reverse complemented and QUAL is stored reversed. An optimisation is conducted for storing the SEQ data: in the common case where the SEQ aligns to the reference genome with no insertions or deletions, and with at most a single mismatch, only the coordinates of the alignment in the reference genome are stored, along with the offset and nature of the single permitted mismatch, if there is one. The compression of the strings prior to storing them in RAM as well as reducing the storage of SEQ strings to a pointer to the reference genome results in manageable RAM usage even for very large BAM and FASTQ files.

Module 4 is run during FASTQ decompression. When reconstructing a FASTQ read, if Module 2 represented any of the read name, SEQ or QUAL components as a reference to a *place* in the BAM file, the information stored by Module 3 for this *place* and this component is used to reconstruct the component in the FASTQ file.

Limitation for paleogenomics data compression: the Deep method will not work well if the BAM data contains alignments of reads generated by collapsing the original R1 and R2 reads to a single read, as is common in ancient DNA applications^15^, while the FASTQ file contains the original, uncollapsed, reads.

## Results

We tested Genozip Deep co-compression with four different publicly available datasets representing a range of experiment types, sequencer technologies and aligners: 1) whole genome sequencing data^8^ sequenced on Illumina HiSeq 2000 and aligned with bwa^9^, and three datasets from the ENCODE portal^10^: 2) whole genome sequencing data sequenced on Oxford Nanopore MinION and aligned with ngmlr^11^; 3) RNA-seq data sequenced on Pacific Biosciences Sequel II and aligned with minimap2^12^; and 4) single-cell RNA-seq data sequenced on Illumina NovaSeq 6000 and aligned with STAR^13^. A list of the ENCODE identifiers, details of data preparation and command line options used can be found in Table S1. We compared compressing these datasets with the Deep method to two other alternative methods. The first method used cutting edge open source tools: we compressed the BAM data into CRAM using samtools^14^ and compressed FASTQ using Spring^1,14^, selected for being the most widely cited FASTQ compression tool. The second method compressed the BAM and FASTQ data, separately, with Genozip. All tools were run in their default compression mode, with command line options indicating the data type when needed: `--long` was specified in Spring for datasets 2 and 3 to indicate long reads and `--pair` was specified in Genozip (without Deep) for dataset 1 to indicate paired-end data. A suitable reference file was provided to Genozip and samtools.

We observe that Genozip with Deep co-compression compressed the four datasets to 24%, 25%, 4.3% and 13% of their original sizes, respectively (Figure 2, Table S2). Note that the original files were already compressed—the BAM files are compressed internally with BGZF and all FASTQ files in these datasets were all in.fastq.gz (gzip) format. We further observe that Deep compression of the four datasets resulted in file sizes smaller than regular Genozip by a factor ranging from 1.9 to 5.7, and smaller than the CRAM/Spring combination by a factor ranging from 2.3 to 6.8 (Figure 2, Table S2).

**Figure 2:**
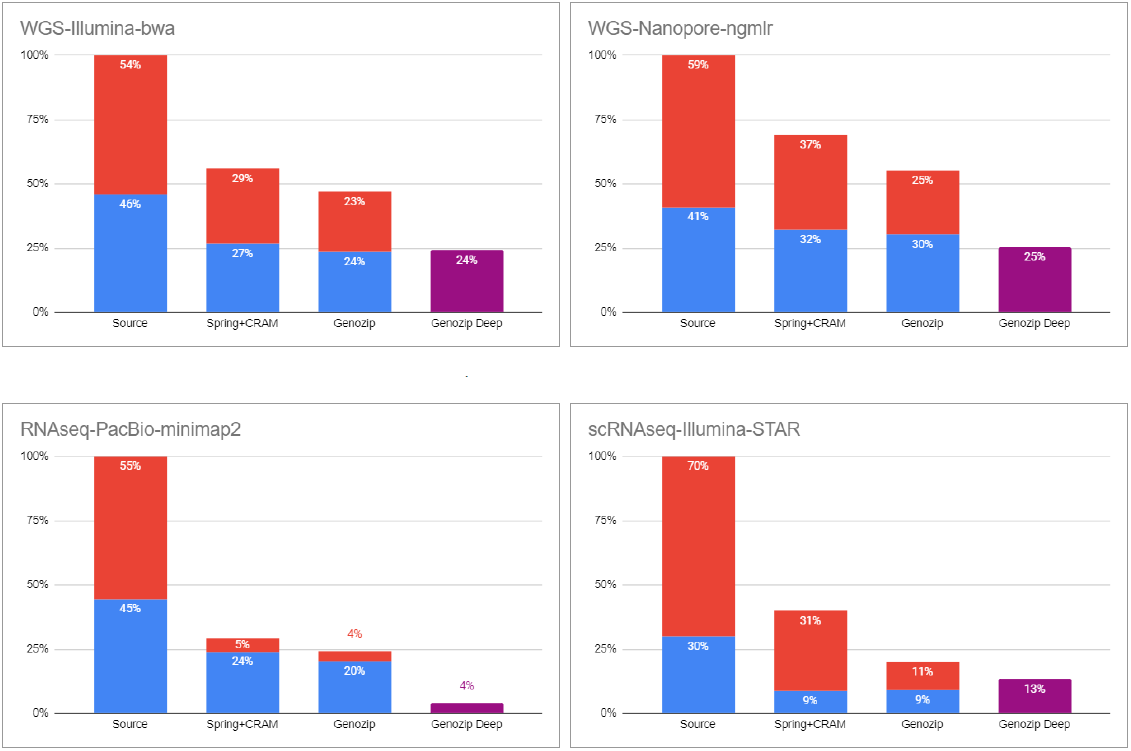
compression comparison. Upper left: whole genome sequenced with Illumina and aligned with bwa (1 BAM file and 2 FASTQ.gz files). Upper right: whole genome sequenced with Oxford Nanopore Technology and aligned with ngmlr (1 BAM and 1 FASTQ.gz file). Bottom left: RNAseq dataset sequenced with Pacific Biosciences and aligned with minimap2 (1 BAM and 1 FASTQ.gz file). Bottom right: single-cell RNA-seq dataset, sequenced with Illumina and aligned with STAR (1 BAM file and 2 FASTQ.gz files). In each panel, the leftmost bar is the original dataset and the other bars represent the three compression methods: Spring^1^ (for FASTQ) + CRAM (for BAM); Genozip; and Genozip Deep. The bars are scaled so that 100% represents the total size of the original dataset. The blue sub-bars represent the relative sizes of the FASTQ data (in case of multiple FASTQ files, their combined size) and the red sub-bars represent the relative sizes of the BAM data. For Deep compression, the resulting file is the co-compression of the entire dataset and is represented in purple.

We ran our tests on a computer with 56 cores. Genozip over-subscribes threads to available cores, resulting in using 64 compute threads. For a fair comparison, we set the number of threads to 64 in samtools and Spring as well. Genozip Deep compressed the 4 data sets in 53, 46, 0.25 and 14.4 minutes, respectively (rounded to two significant digits), which is a bit faster than the 57, 52, 0.4 and 14.6 minutes consumed by regular Genozip and significantly faster than the 149, 88, 1.3, 269 minutes consumed by the CRAM/Spring combination. More details on compression times can be found in Table S3. Decompression of a Genozip Deep file took 37, 36, 0.33 and 10 minutes, respectively, which is mostly marginally better than regular Genozip with 42, 39, 0.32 and 11 minutes, and roughly similar to the CRAM/Spring combination with 31, 38, 0.85 and 10 minutes. More details on decompression times are in Table S4.

Genozip Deep method has a drawback related to its RAM consumption. When compressing the four datasets, the maximum physical RAM usage reached 115 GB, 132 GB, 9 GB, and 95 GB, respectively. This consumption is higher than for the other methods, with regular Genozip utilising 52 GB, 130 GB, 8 GB, and 82 GB, and the CRAM/Spring combination using 40 GB, 87 GB, 9 GB, and 14 GB, respectively. Further information on memory consumption can be found in Table S3, specifically under the “maximum resident set” category. Genozip is designed to liberally use as much RAM as it requires to maximise compression. However, the user may modify this default behaviour with the `--low-memory` command line option, which directs Genozip to conserve RAM even at the expense of the compression ratio.

In conclusion, Genozip Deep addresses the common need for long-term archival of FASTQ and related BAM files with the best available compression, significantly better than other current best-performing solutions.

## Supporting information

Benchmark raw data

## Notes

https://www.genozip.com

### Competing Interest Statement

1) This research is funded by licensing fees paid by commercial users of the Genozip to Genozip Limited, a company wholly owned by DL.
2) The Deep method is patent pending, USPTO application # 18204466
3) Institute for Genomic Medicine, Columbia University is a paying customer of Genozip.

